# Prognostic and biological roles of Parkinson’s disease genes in cancer

**DOI:** 10.1101/2025.03.22.644759

**Authors:** Sara Fiume, Francesca Molinari, Benedetta Vai, Sara Poletti, Giovanni Citterio, Ferdinando Fiumara, Stephanie Papin, Paolo Paganetti, Maurizio Callari, Luca Colnaghi

**Affiliations:** Division of Neuroscience, IRCCS San Raffaele Scientific Institute, Milan, Italy; IRCCS San Raffaele Scientific Institute, Milan, Italy; Rita Levi Montalcini Department of Neuroscience, University of Torino, Corso Raffaello 30, Torino 10125, Italy; Institute of Oncology Research (IOR), CH6500 Bellinzona, Switzerland; Faculty of Biomedical Sciences, Università della Svizzera Italiana, Lugano, Switzerland; Fondazione Michelangelo, Milan, Italy; School of Medicine, Vita-Salute San Raffaele University, Milan, Italy

## Abstract

**Background:** Increasing evidence suggests significant associations between Parkinson disease (PD) and cancer risks, with epidemiological studies revealing a complex relationship. PD patients exhibit lower risks of lung, genitourinary, and gastrointestinal cancers but higher risks of melanoma and brain cancers. Despite these observations, the underlying mechanisms between PD and cancers are poorly understood.

**Objectives:** We aimed to identify molecular signatures that could provide insight into this complex connection by assessing the association of PD-related genes with patient survival and the cancer-specific co-expression networks they are involved in.

**Methods:** To explore this, we analyzed transcriptomic data from 18 cancer types in the TCGA dataset (n=6,088) and 16 cancer types in the DepMap dataset (n=682). We focused on seven genes causally implicated in PD (*SNCA*, *PINK1*, *LRRK2*, *PRKN/PARK2*, *PARK7*, *GBA1*, and *ATP13A2*) and conducted *in silico* analyses, to evaluate their associations with survival and correlation with genes, pathways and response to drugs in the context of cancer.

**Results:** Our findings revealed that the expression levels of the genes correlated with overall survival in a cancer-specific manner, often influenced by the TP53 genetic status. These genes were also associated with key cancer hallmarks such as genomic instability and cell proliferation. Additionally, novel associations were identified linking these genes to drug responses in a context-specific manner.

**Conclusions:** This study suggests that PD and cancer may be linked by biological pathways, sometimes associated with cancer hallmarks. These findings provide potential insights into druggable targets and the shared molecular mechanisms underlying PD and cancer.

## Introduction

Parkinson’s disease (PD) is a progressive neurodegenerative disorder primarily characterized by motor symptoms such as tremors, bradykinesia, rigidity, and postural instability, as well as a range of non-motor symptoms including cognitive impairment, mood disorders, and autonomic dysfunction (1). Traditionally, PD has been associated with the degeneration of dopaminergic neurons in the substantia nigra (2), but emerging evidence suggests that it could also be linked to other pathological processes that might influence the risk of other diseases, including cancer (3).

Epidemiological studies investigating the link between PD and cancer generally indicate an inverse relationship (4). Individuals with PD tend to have a lower overall risk of developing cancer, and cancer patients have a reduced risk of developing PD. This inverse association could be explained by a lower incidence of smoking-related cancers (such as lung, bladder, and colorectal cancers) among individuals with PD, likely due to the lower prevalence of smoking within this population (5). However, positive correlations between PD and certain cancers, including melanoma, skin, breast, brain, and prostate cancers, have also been documented (4,6 and 7), sometimes with confounding results (8). Among these possible correlations, the positive association between PD and melanoma seems to be the most established (9). This link is observed for melanomas developing both prior to and following a PD diagnosis, suggesting that antiparkinsonian medications are unlikely to be the underlying cause (10). Common genetic factors could explain this association, as people with a familial history of melanoma had an increased risk for PD. This was recently confirmed by using GWAS data from two PD and six cancer consortia (11).

Most PD cases are sporadic, with only about 10-15% having a positive family history (12). The most significant genetic risk factors for PD are mutations in the *GBA1* gene, which encodes the lysosomal enzyme glucocerebrosidase, essential for glycosphingolipid homeostasis (13). Clinically, PD associated with *GBA1* mutations mirrors sporadic PD but typically manifests earlier, with a younger age at onset, more frequent cognitive impairment, and a faster disease progression (13). At least other six genes have been firmly linked to hereditary monogenic PD. Mutations in *SNCA* (*PARK1/4*, Alpha-Synuclein) and *LRRK2* (*PARK8*, Leucine-Rich Repeat Kinase 2) cause autosomal-dominant PD.

Meanwhile, mutations in *PRKN* (*PARK2*, Parkin), *PINK1* (*PARK6*, PTEN-Induced Putative Kinase 1), *PARK7* (DJ-1, Parkinsonism Associated Deglycase), and *ATP13A2* (*PARK9*, ATPase Type 13A2) lead to autosomal recessive forms (14). These genetic distinctions underscore the complex inheritance patterns and molecular pathways contributing to PD.

Interestingly, most of these genes have also been implicated in molecular mechanisms traditionally associated with cancer. For instance, *SNCA*, *LRKK2*, *PRKN/PARK2*, *PINK1*, and *PARK7* have been reported to play roles in the DNA damage response and the maintenance of genomic stability (15–19). *PINK1* has been shown to interact with *PTEN*, a gene frequently mutated in various cancers (20), and *SNCA* has been found to functionally interact with *TP53* (21). All these associations raise intriguing questions about the underlying biological mechanisms that may link PD with cancer susceptibility.

To further explore this connection, it is essential not only to investigate the genetic roots of PD but also to shed light on the molecular mechanisms that may link PD genes to cancer biology.

By leveraging data from The Cancer Genome Atlas (TCGA) (22) and the DepMap datasets (23), this study aims to elucidate the expression patterns of these PD-associated genes across different cancer types, assess their association with patient survival, and unveil the biological co-expression networks they are involved in. Understanding these patterns may reveal novel insights into the bidirectional relationship between PD and cancer, shedding light on shared pathophysiological pathways and identifying potential targets for therapeutic intervention.

## Methods

### Datasets

#### TCGA Dataset analysis

The Cancer Genome Atlas (TCGA) pan-cancer dataset was obtained from the GDC portal, specifically utilizing the files EBPlusPlusAdjustPANCAN_IlluminaHiSeq_RNASeqV2.geneExp.tsv and clinical_PANCAN_patient_with_followup.tsv. This dataset initially comprised 11,069 samples spanning 32 distinct cancer types. After excluding 737 non-tumoral samples, the final cohort for analysis consisted of 10,332 samples.

For downstream analyses, genes were retained if they exhibited an average expression level of log10(FPKM + 1) greater than 0.5 and a standard deviation exceeding 0.2 in at least one cancer type, resulting in a filtered set of 17,646 genes.

The expression levels of *SNCA, PINK1, LRRK2, PRKN/PARK2, PARK7, GBA1*, and *ATP13A2* were correlated with those of all other genes using Spearman correlation, as implemented in the stats package (v. 3.5.0) in R.

Additionally, *TP53* mutation and copy number alteration (CNA) data, also downloaded from the GDC portal, were utilized to classify samples into wild-type (WT) and mutant (MUT) groups, as well as to delineate functionally distinct subgroups according to the criteria outlined in Callari et al. (24).

#### DepMap dataset analysis

MAPT CRISPR KO, gene expression, proteomic, and drug response data from pan-cancer cell lines were downloaded from the Dependency Map initiative (https://depmap.org/portal/). Specifically, we obtained the following files: from version 21Q1 – CCLE_expression.tsv, protein_quant_current_normalized.csv, sanger-dose-response.csv, and sample_info.csv; and from PRISM Repurposing 19Q4 – primary-screen-replicate-collapsed-logfold-change.csv. Only cancer types represented by more than one cell line in the gene expression dataset were retained, and cell lines annotated as “Engineered” or “Fibroblast” were excluded. After these filters, gene expression data were available for 1,320 cell lines across 28 cancer types, while drug response data were available for 692 cell lines representing 22 cancer types.

For each cancer type, the area under the drug response curve (AUC)—which ranges from 0 (indicative of response) to 1 (indicative of resistance)—was correlated with *SNCA, PINK1, LRRK2, PRKN/PARK2, PARK7, GBA1,* and *ATP13A2* expression.

Spearman correlation analysis was conducted only when a cancer type included at least 10 cell lines and contained a minimum of two cell lines classified as sensitive (AUC < 0.8) and two as resistant (AUC > 0.8).

### Data analysis

#### Correlation analysis

All statistical analyses were conducted in R (v 4.4.1; platform: x86_64-apple-darwin17.0, macOS Monterey 12.3). Associations between pairs of continuous variables (e.g., gene expression levels) were quantified using Spearman’s correlation analysis. P-values were computed with the *cor.test* function and adjusted for multiple comparisons using the Benjamini-Hochberg correction using the *p.adjust* function.

#### Unsupervised analysis

Cancer types were clustered based on the correlation profiles of *SNCA, PINK1, LRRK2, PRKN/PARK2, PARK7, GBA1,* and *ATP13A2* expression. Clustering was performed using the ConsensusClusterPlus method (25), implementing the K-means algorithm with Euclidean distance as the similarity metric.

#### GeneSet Enrichment Analysis (GSEA)

A custom list of gene sets was assembled as follows. The HALLMARK gene set collection (v7.1) was obtained from MSigDB (26), and gene sets related to the PANTHER classification system (27) were downloaded from https://maayanlab.cloud/Harmonizome/dataset/PANTHER+Pathways.

Correlation-ranked genes were then subjected to gene set enrichment analysis using the gsea function from the phenoTest R/Bioconductor package (v1.28.0). Gene sets with a false discovery rate (FDR) below 0.1% and an absolute normalized enrichment score (NES) greater than 2.3 in at least one cancer type were considered significant and reported. Additionally, in analyses stratified by *TP53* status, gene sets displaying an absolute delta NES (i.e., the difference in NES between *TP53* mutant and wild-type tumors) exceeding 0.6 were deemed significant.

#### Survival analysis

Overall survival was evaluated using univariate and multivariate Cox regression (28–29) analyses implemented in the survival (v3.1) R/Bioconductor package. Analyses were performed only when at least 10 events (deaths) were observed within a given cancer case subset. For multivariate models, at least three of the following four covariates— tumor size, lymph node status, metastatic status, and AURKA gene expression—needed to be available in a minimum of 20 patients. A significance threshold of p < 0.05 was applied.

Gene expression levels for *SNCA, PINK1, LRRK2, PRKN/PARK2, PARK7, GBA1,* and *ATP13A2* were dichotomized into high and low groups using the median expression within each cancer type as the cutoff.

Hazard ratios (HRs) were calculated from the Cox regression coefficients to quantify the relative risk associated with gene expression. An HR below 1 indicates a decreased risk relative to the reference group, an HR above 1 indicates an increased risk, and an HR of 1 denotes no difference in risk.

## Results

### Expression patterns of Parkinson’s disease genes in cancer

To elucidate the relevance of PD genes in cancer, we began by evaluating the expression of 7 Parkinson’s disease genes (*SNCA, PINK1, LRRK2, PRKN/PARK2, PARK7, GBA1,* and *ATP13A2)* across 18 distinct cancer types using transcriptomic data from the TCGA pan-cancer cohort. The cancers were selected from the available 32 to represent lung, genitourinary, gastrointestinal, melanoma, and brain tumors. Our analysis revealed substantial variability in gene expression levels across different tumor types, as shown in Figure 1a. Notably, certain cancers, including glioblastoma multiforme (GBM), lower grade glioma (LGG), uveal melanoma (UVM), and skin cutaneous melanoma (SKCM), consistently exhibited higher expression levels of the PD-related genes. This was particularly evident for *SNCA, ATP13A2,* and *GBA1,* whose elevated expression suggests potential involvement in tumor biology or stress responses unique to these malignancies. Interestingly, not all PD-related genes followed the same expression patterns. For instance, *LRRK2* displayed peak expression in prostate adenocarcinoma (PRAD) and lung adenocarcinoma (LUAD), suggesting its role may extend to cellular pathways active in epithelial cancers. *PARK7* showed the highest expression in diffuse large B-cell lymphoma (DLBC) and LUAD. In contrast, cancers such as esophageal carcinoma (ESCA) and colon adenocarcinoma (COAD) demonstrated relatively low expression levels for most of the genes, potentially reflecting a less pronounced role of PD-associated pathways in these tumor types. We next compared the expression of the genes to each other and found that unsupervised clustering of their correlation patterns identified distinct cancer groups (Figure 1b). Notably, LGG, GBM, SKCM, and UVM formed a cluster characterized by higher expression levels of these genes. These findings, in conjunction with epidemiological evidence, suggest a potential role for PD-related genes in these cancers, warranting further investigation into their tumor-specific functions.

**Figure 1:**
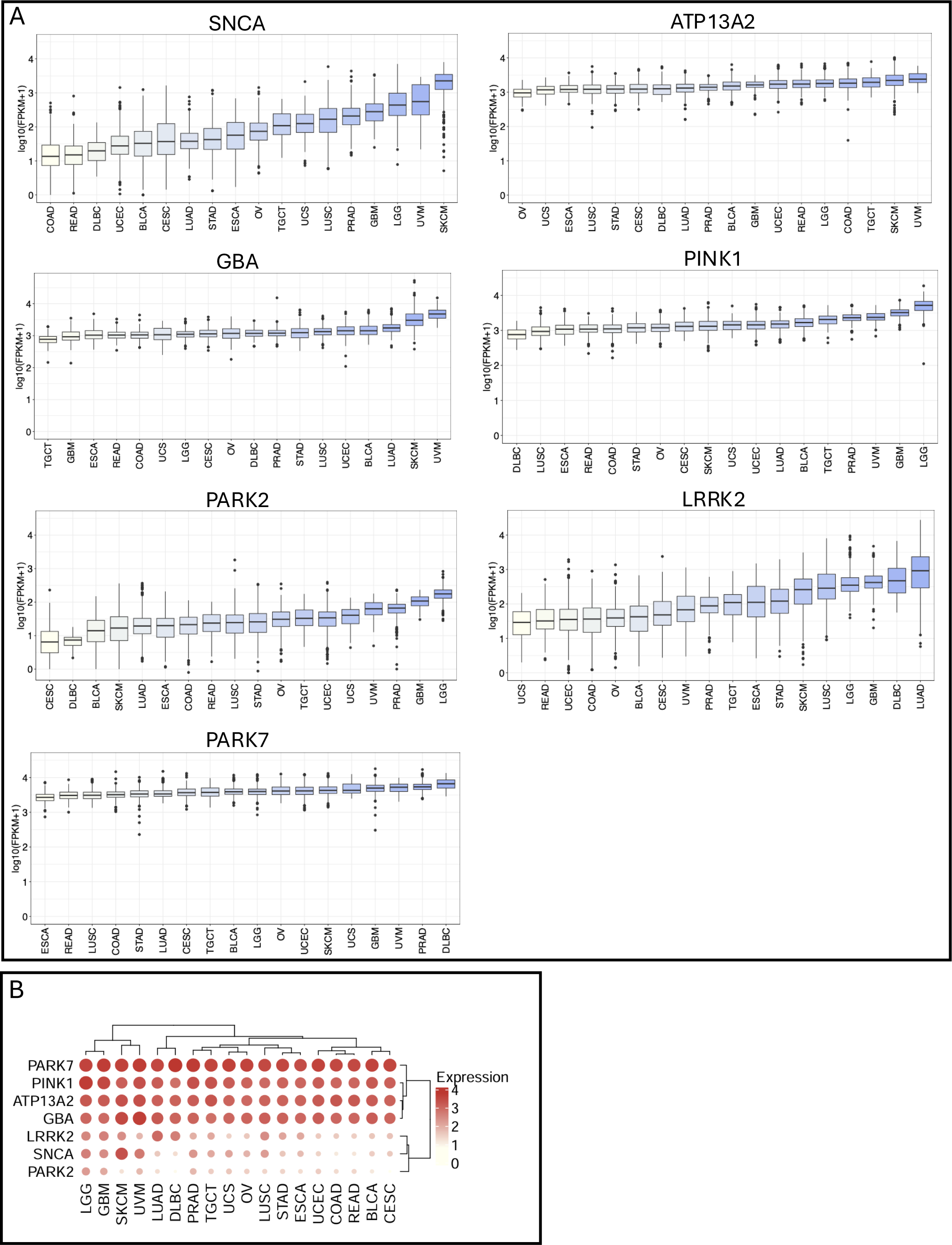
A) Pan-cancer evaluation of PD-related genes’ expression and transcriptional associations obtained by mining the TCGA dataset. B) Correlation expression among SNCA, PINK1, LRRK2, *PRKN/PARK2*, PARK7, GBA1, and ATP13A2.

### PD gene association with survival in skin and brain cancers

Considering the observed expression patterns, we next investigated the connection between the expressions of *SNCA, PINK1, LRRK2, PRKN/PARK2, PARK7, GBA1,* and *ATP13A2* and overall survival (OS) in UVM, SKCM, GBM, and LGG. First, we applied univariate Cox regression analysis for each cancer in the overall population. Next, we performed multivariate Cox regression analysis, adjusting for tumor size, lymph node status, metastatic status, and expression of the AURKA gene, which serves as a marker of the proliferation process, associated with survival in multiple cancers (Table 1).

**Table 1:**
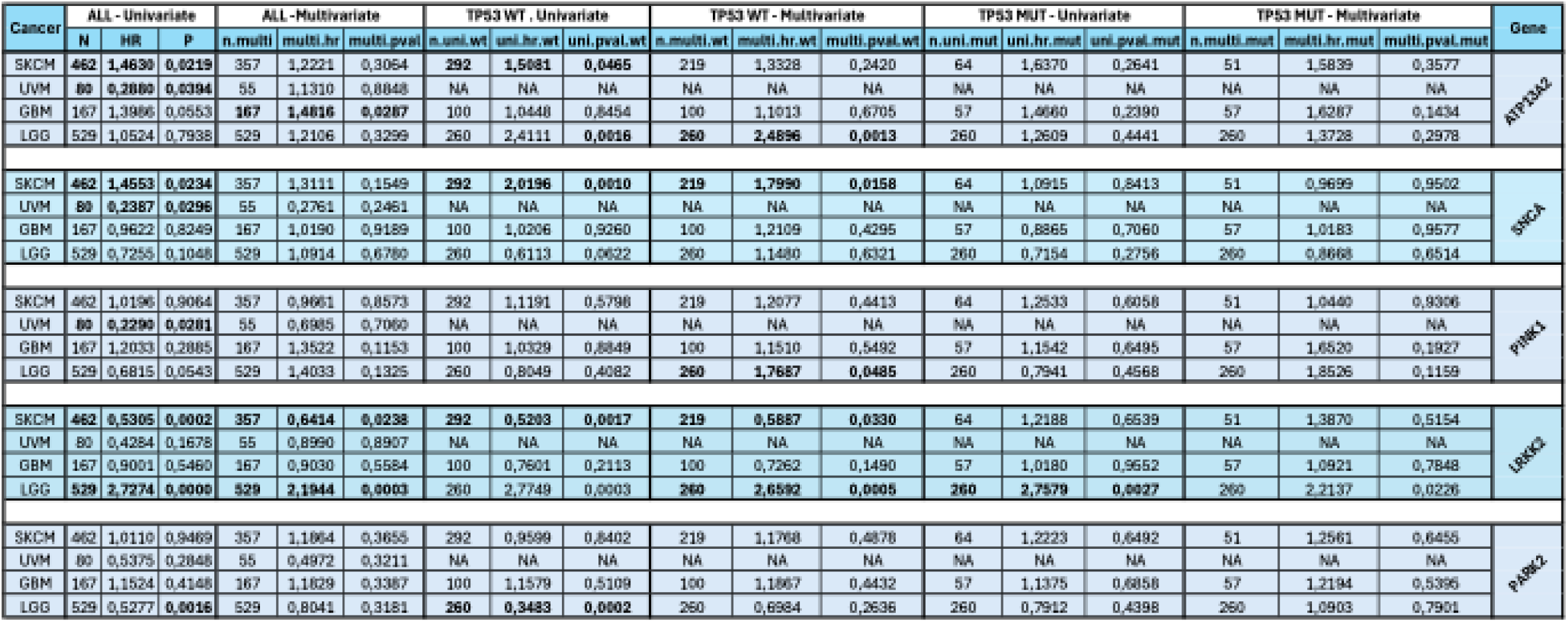
Univariate and multivariate Cox survival analysis for each cancer type in all samples and stratified by P53 status.

Kaplan-Meier survival curves were generated to visually depict whether gene expression levels correlated with patient outcomes. In UVM, the expressions of *SNCA, PINK1,* and *ATP13A2* demonstrated significant correlations with survival outcomes (Figure 2a). Higher expression of these genes was associated with improved OS, suggesting potential protective or adaptive roles in this cancer type. These findings suggest that elevated expression of these PD-related genes might counteract tumor progression or contribute to cellular resilience in UVM (Figure 2a). We did not find significant differences for the other four genes. In SKCM, gene-specific survival correlations reflected both positive and negative associations (Figure 2b). *LRRK2* expression was positively correlated with survival, as patients with higher expression exhibited better OS (p= 1e-4). Conversely, *SNCA* and *ATP13A2* displayed negative correlations with survival, where higher expression levels were linked to poorer outcomes (p= 0.02 for both) (Figure 2b). We did not find significant difference for the other genes.

**Figure 2:**
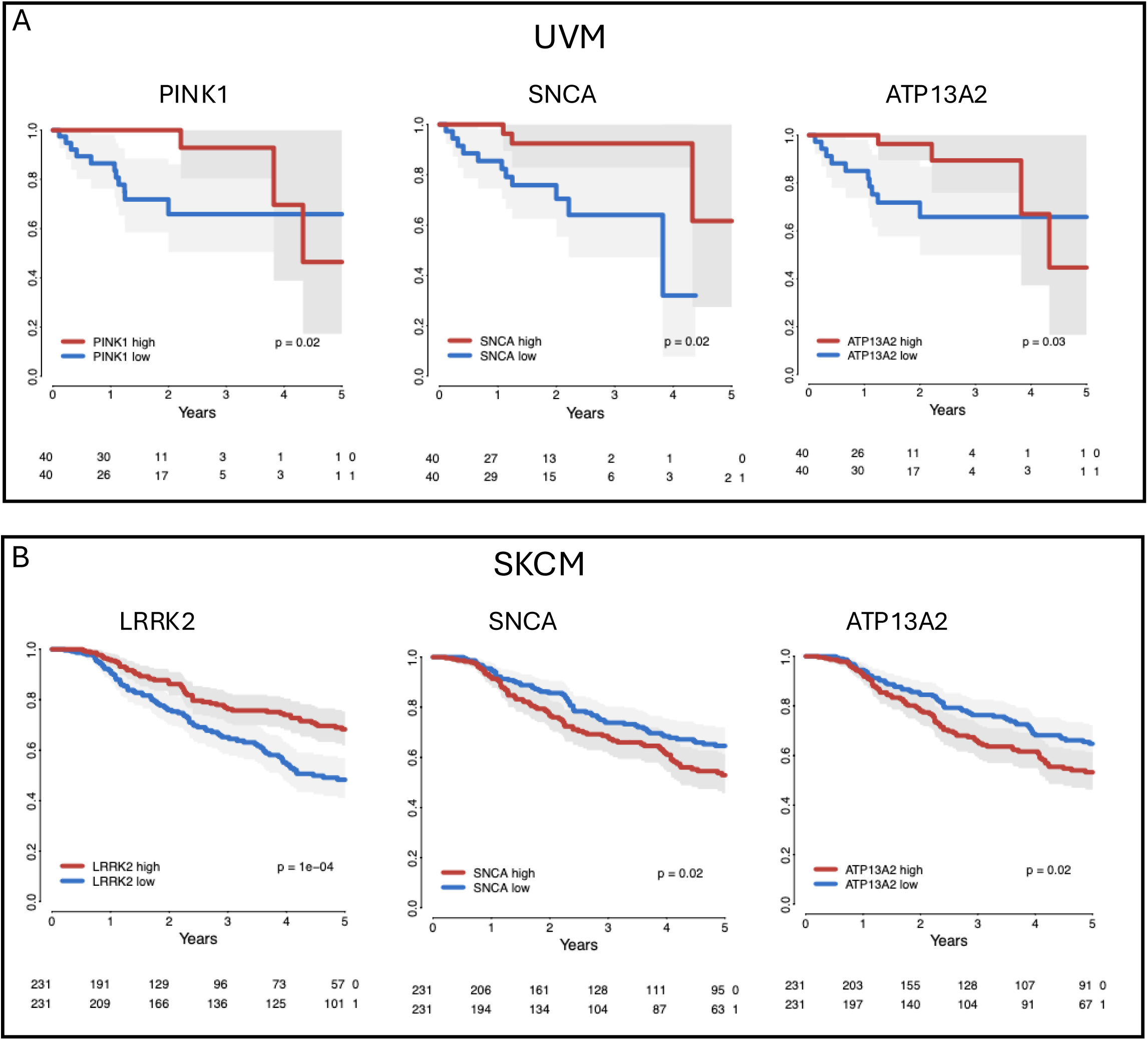
A-B) Selected Kaplan-Meier curves showing the association between PD-related genes and survival in UVM and SKCM cancer types. P-value obtained by log-rank test.

**Figure 3:**
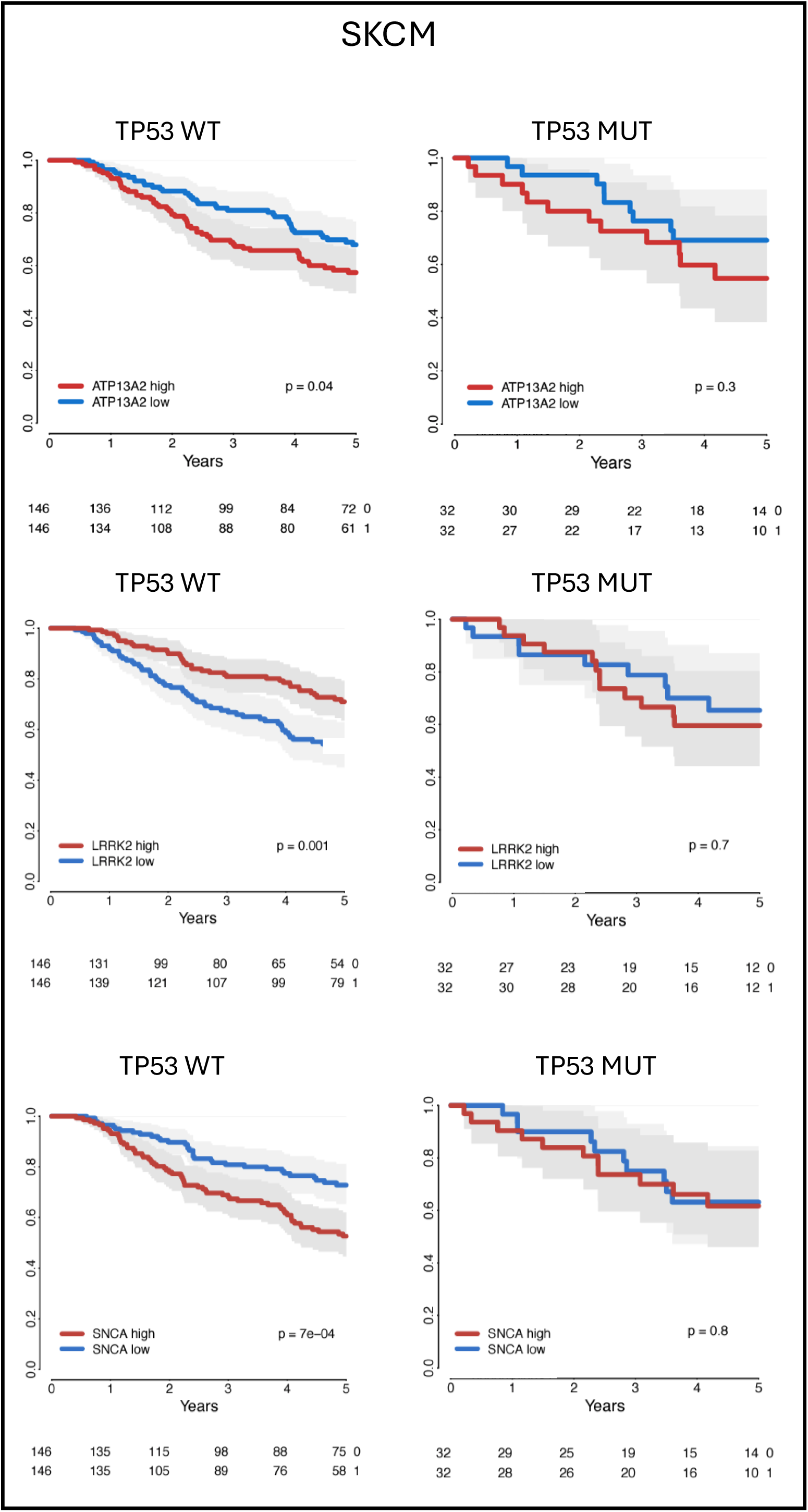
Selected Kaplan-Meier curves showing the association between PD-related genes and survival in SKCM, stratified by TP53 status. P-value obtained by log-rank test.

Given the critical role of *TP53* in tumor biology (30), we stratified the survival analysis by *TP53* status (wild type [WT] or mutated [MUT]) to assess whether the prognostic value of *SNCA, LRRK2*, and *ATP13A2* depends on *TP53* functionality. In UVM, stratification by *TP53* status did not reveal significant survival differences for *SNCA, LRRK2*, or *ATP13A2*. In SKCM, stratification by *TP53* status uncovered patterns of association with survival for *SNCA, LRRK2*, and *ATP13A2*, specifically in the *TP53* WT subset. High expression levels of *ATP13A2* significantly correlated with improved OS in *TP53* WT SKCM (p= 0.04). However, no significant association was observed in the *TP53* MUT subset (p= 0.3). High expression of *LRRK2* was strongly associated with better survival in the *TP53* WT cohort (p= 0.001) but again not in the *TP53* MUT subset (p= 0.7). In contrast, high expression of *SNCA* was negatively correlated with OS in *TP53* WT SKCM (p= 7e-4). However, no significant association was observed in the *TP53* MUT group (p= 0.8).

We next studied the correlations in brain tumors. In LGG, patients with higher levels of *PINK1* expression exhibited a trend toward improved OS compared to those with lower *PINK1* expression, with a p-value of 0.05 (Figure 4a). High *LRRK2* expression was associated with significantly better OS than low expression, with a highly significant p-value of 1e-06. Similarly, *PRKN/PARK2* expression also demonstrated a significant impact on survival. Patients with higher *PRKN/PARK2* levels showed better OS than those with lower levels (p= 0.001). Next, we stratified for *TP53* status. In LGG with *TP53* WT, high *PRKN/PARK2* expression was significantly associated with better OS (p= 1e-04). However, this relationship was not observed in LGG with *TP53* mutations. These results suggest that *PRKN/PARK2’s* prognostic value may depend on an intact *TP53* pathway. Similarly, *LRRK2* expression demonstrated a strong correlation with survival in LGG with *TP53* WT. Patients with high *LRRK2* levels showed improved OS compared to those with low *LRRK2* expression (p= 1e-04), but this was lost in the *TP53* mutant subgroup (p= 0.4). This pattern was found also for *ATP13A2* expression, which was associated with better OS in LGG patients with *TP53* WT (p= 0.001) but not in the *TP53* mutant context (p= 0.4) (Figure 4a). Interestingly, in GBM, PARK7 expression was associated with both survival and TP53 status (Figure 4b). Among TP53 wild-type (WT) cases, high PARK7 expression correlated with slightly improved survival outcomes compared to low expression (p= 0.02). However, in TP53-mutant GBM, no significant difference in survival was observed between high and low PARK7 expression groups (p= 0.5) (Figure 4b).

**Figure 4:**
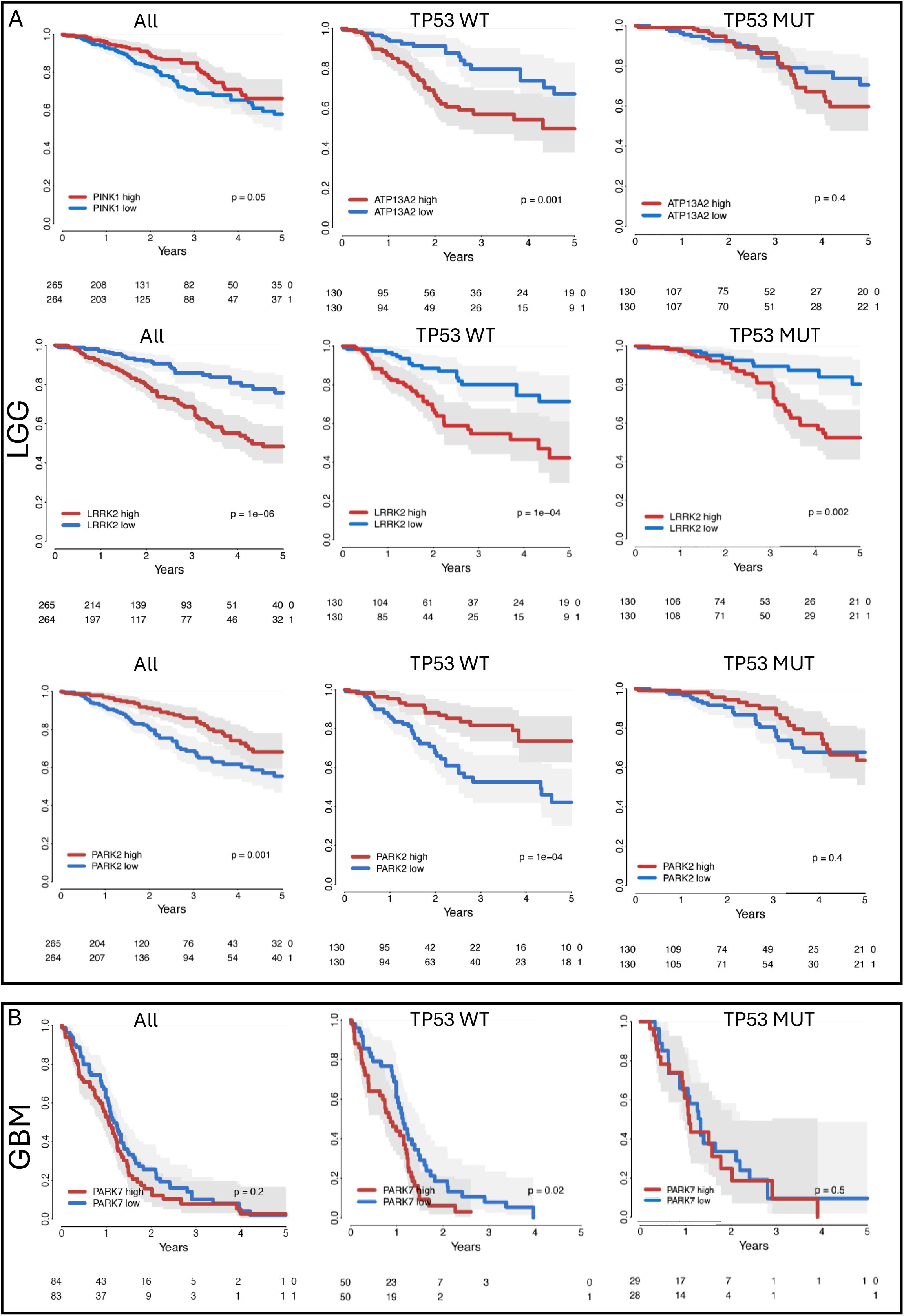
A) Selected Kaplan-Meier curves showing the association between PD-related genes and survival in LGG. P-value obtained by log-rank test. B) Selected Kaplan-Meier curves showing the association between PD-related genes and survival in LGG and GBM, stratified by TP53 status. P-value obtained by log-rank test.

Examining the Hazard Ratio presented in Table 1, *ATP13A2* expression in SKCM showed a significant association with survival in both univariate (hazard ratio [HR]= 1.463, p= 0.0219) and multivariate analyses (HR= 1.222, p= 0.3064). When stratified by *TP53* status, *ATP13A2* demonstrated significant prognostic value in the *TP53* WT subgroup (HR= 1.7990, p= 0.0158) but not in the *TP53* mutant subgroup. Similarly, in LGG, *ATP13A2* expression was significantly correlated with survival in the overall population and *TP53* WT subgroup (HR= 2.4896, p= 0.0013), with no significant association observed in the *TP53* mutant subgroup. *SNCA* expression was significant only in UVM, where higher expression was associated with better survival in the univariate analysis (HR= 0.229, p= 0.0281). However, this relationship was not maintained in multivariate analysis or when stratified by *TP53* status. *PINK1* expression was significantly associated with survival in LGG in the *TP53* WT subgroup (HR= 1.7687, p= 0.0485). Although *PINK1* demonstrated no significant association in other cancers or in the *TP53* mutant subgroup, these findings highlight its potential prognostic importance in LGG with intact *TP53* pathways. In the overall population, *LRRK2* showed a significant positive correlation with survival (HR= 2.7274, p= 0.0002). This effect was particularly pronounced in the *TP53* WT subgroup, where multivariate analysis revealed an HR of 2.6592 (p= 0.0005). Interestingly, *LRRK2* also demonstrated significance in the *TP53* mutant subgroup (HR= 2.2137, p= 0.0226), indicating its potential as a prognostic marker independent of *TP53* status in LGG. *PRKN* showed significant prognostic value in LGG in both univariate (HR= 2.7274, p= 0.0002) and multivariate analyses (HR= 2.8041, p= 0.0003). While the *TP53* WT subgroup demonstrated a strong association (HR= 3.3483, p= 0.0002), no significant correlation was observed in the *TP53* mutant subgroup. Overall, this analysis suggests that *PINK1, SNCA, LRRK2,* and *ATP13A2* may be the PD-related genes with the greatest influence on UVM, SKCM, GBM, and LGG.

### Gene and pathway correlation analysis for PD genes in skin and brain cancer

To further explore the molecular mechanisms underlying these associations, we examined the correlation between the expression levels of *PINK1, SNCA, LRRK2,* and *ATP13A2* and all expressed genes in the four cancer types (UVM, SKCM, LGG, and GBM). The ranked gene lists were then used as input for Gene Set Enrichment Analysis (GSEA) to identify biological processes and pathways either positively or negatively associated with their expression. We found that *PINK1, SNCA, LRRK2,* and *ATP13A2* expression levels were positively or negatively correlating with several cancer pathways, including those related to cell cycle regulation, apoptosis, and DNA damage response (Figure 5). The PD-associated genes negatively correlated with cell cycle and proliferation genesets, particularly the G2M checkpoint and E2F targets. Pathways related to oxidative phosphorylation and fatty acid metabolism were notably enriched, suggesting a potential link between PD-related mitochondrial dysfunction and cancer metabolism, potentially impacting tumor growth and survival through altered energy metabolism. Additionally, PD-associated genes were enriched in pathways involving apoptosis and DNA repair mechanisms, indicating roles in maintaining genomic integrity and potentially contributing to cancer cell survival and resistance to therapy through the dysregulation of apoptosis pathways. The enrichment of inflammatory response pathways, including interferon alpha and gamma responses, points to a possible influence of PD genes on cancer development via modulation of the immune response. Furthermore, the pathway enrichment analysis revealed variability across different cancer types, such as GBM, LUSC, and SKCM, highlighting the context-dependent role of PD genes in cancer biology, where they may act as oncogenes or tumor suppressors depending on the context.

**Figure 5:**
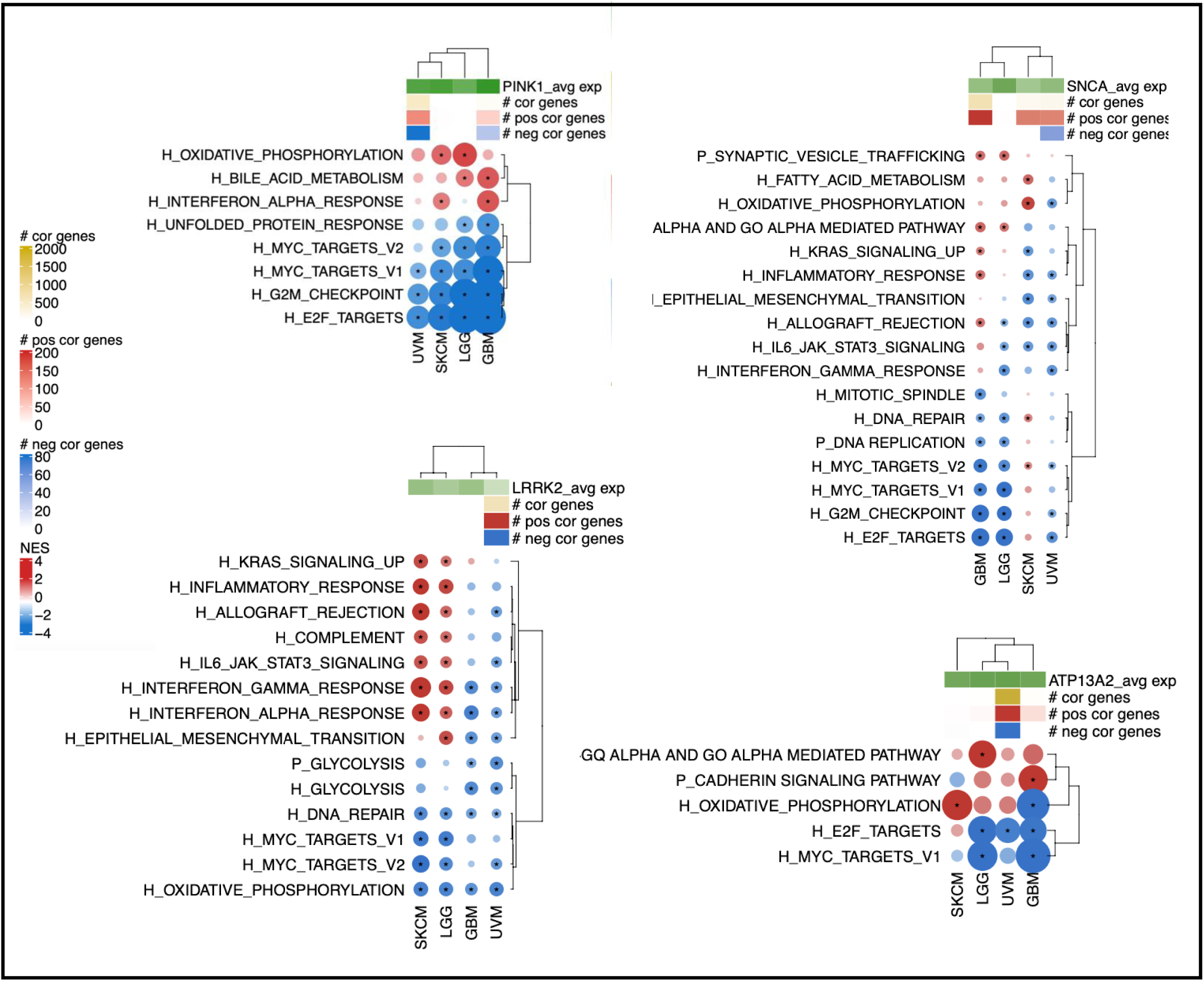
GeneSet Enrichment Analysis on the genes ranked according to their correlation with *PINK1*, *SNCA*, *LRRK2*, and *ATP13A2* expression in each cancer type. A negative Normalised Enrichment Score (NES) means down-regulation of the geneset for high PD related genes’ expression and vice versa for positive NES.

### PD genes can predict response to cancer drugs

Next, leveraging the DEPMAP transcriptomic data collection, we analyzed the expression patterns of *PINK1, SNCA, LRRK2*, and *ATP13A2* in tissue-specific cancer cell lines. This approach provided valuable insights into how these genes are expressed in cellular models of cancer, complementing our findings from the TCGA dataset. The analysis, although revealing distinct expression patterns for key PD-associated genes, confirmed the TCGA finding that expression on average was higher in the nervous system and skin-derived cells (Figure 6a). We next analyzed the CRISPR DEPMAP data collection for the cell viability after *PINK1, SNCA, LRRK2,* and *ATP13A2 knockout (*KO). The impact of the KO on viability was cancer and gene specific. Interestingly, an overall decrease in viability is prevalently observed, except for *SNCA* cell lines (Figure 6b).

**Figure 6:**
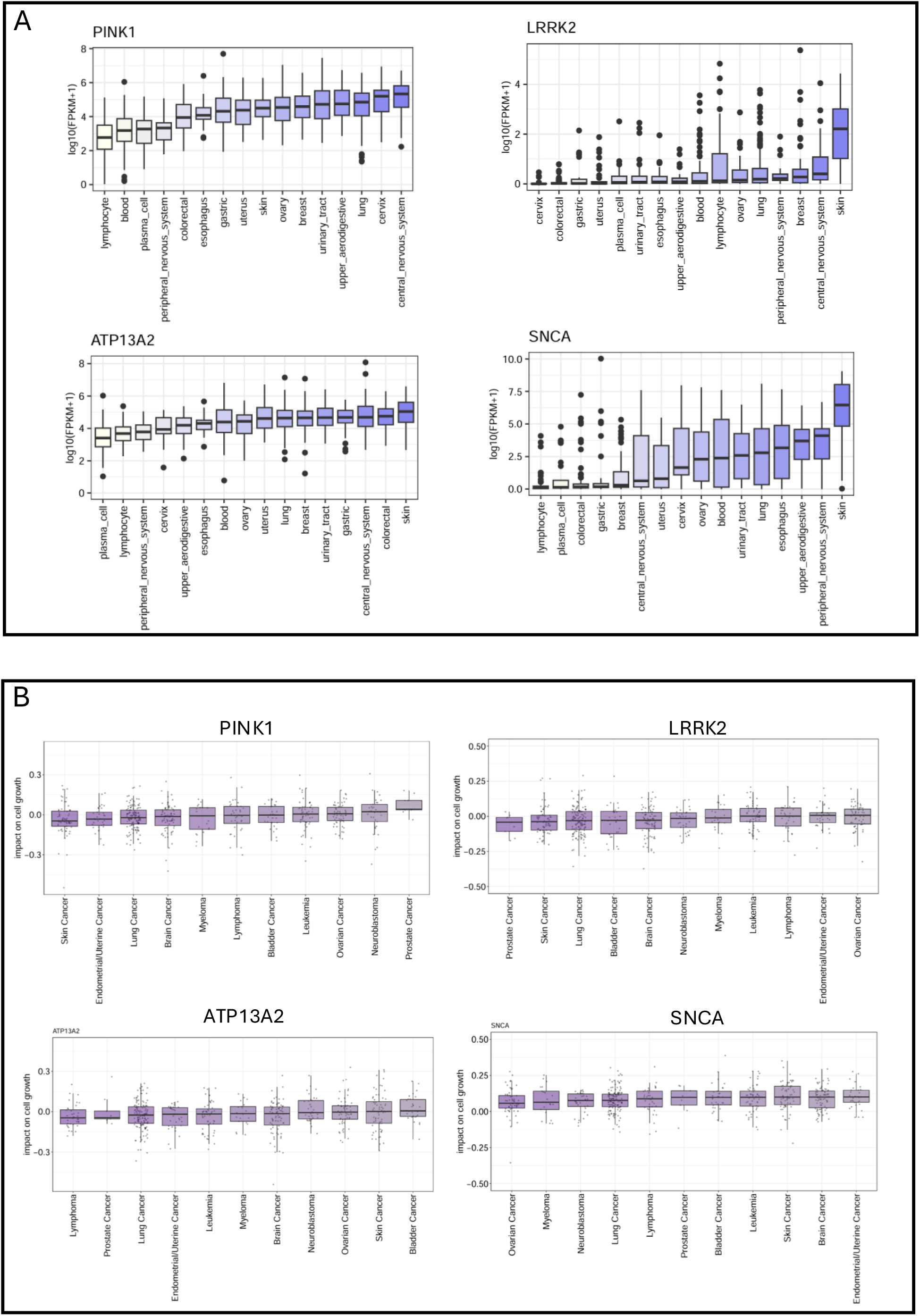
A) PD-related genes’ expression in the DEPMAP cell line dataset according to the cancer type. B) Cell viability after PD-related genes KO in the CRISPR DEPMAP dataset. Viability scores are normalized such that nonessential genes have a median score of 0 and independently identified common essentials have a median score of −1.

To translate these insights into potential therapeutic opportunities, we explored the drug response profiles associated with *PINK1, SNCA, LRRK2,* and *ATP13A2* KO. After data filtering, we evaluated 121 drugs in cell lines from 3 cancer types: skin, central (CNS) and peripheral nervous systems (PNS) (Figure 7). The associations involved multiple drug families and were largely specific to the cancer type. For *SNCA* KO, several compounds had a positive correlation in skin-related cell lines. A different scenario was present for PNS-related cell lines, where compounds had both positive and negative correlations. For the central nervous system, the correlations were less strong.

**Figure 7:**
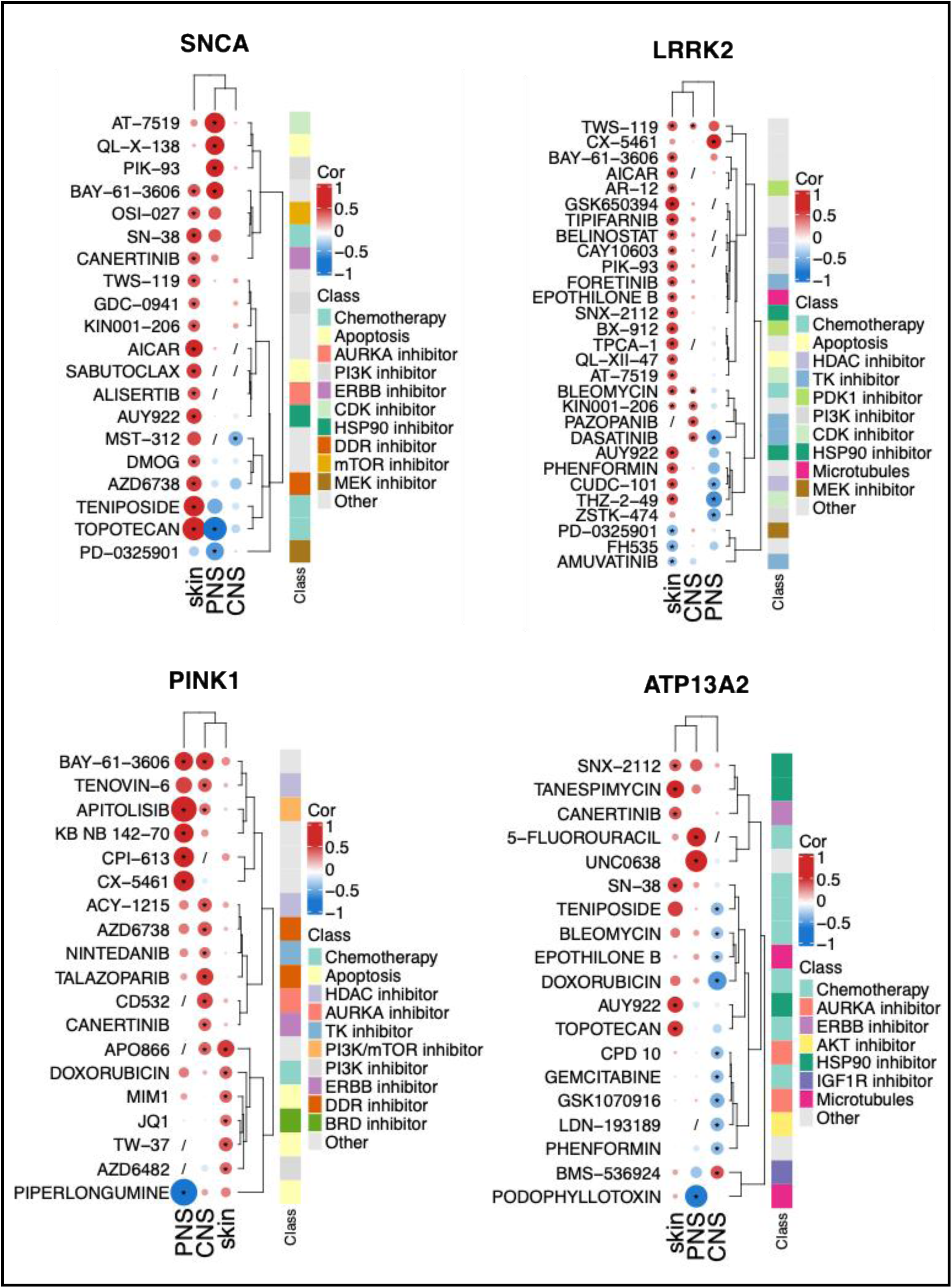
Heatmap summarizing the correlations between PD-related genes expression with drug response quantified as area under the drug response curve (AUC).

*LRRK2 KO* cell line exhibited sensitivity to a range of inhibitors, with significant effects observed in PNS and CNS tissues. *PINK1* KO cell line drug response profiles revealed efficacy with compounds particularly in PNS and skin tissues. *ATP13A2* KO cell line was linked to several chemotherapeutic agents, including SNX-2112 and Tanespimycin, with notable responses in skin and CNS tissues, indicating its potential as a target in cancers affecting these areas. Overall, these findings underscore the potential of PD-related gene expressions as a possible biomarker for predicting drug response in multiple cancer types.

## Discussion

By leveraging publicly available datasets such as the TCGA, we examined expression patterns, survival associations, and interactions with *TP53* status, revealing the multifaceted nature of PD-related gene involvement in cancer. These results underscore the complexity of the bidirectional relationship between neurodegeneration and oncogenesis, offering new directions for research and therapeutic development.

### Prognostic Implications

We found an elevated expression of PD-associated genes in cancers such as GBM, LGG, UVM, and SKCM that have already been linked to PD (31). This high expression may be linked to the functional roles of these PD-associated genes in cancers. Conversely, the low expression of these genes in ESCA and COAD suggests that PD-related pathways may not be universally exploited in tumorigenesis. This disparity highlights the influence of the tissue microenvironment and tumor-specific genetic drivers in determining the relevance of these genes.

Our survival analyses revealed that PD-related gene expression has distinct prognostic implications across cancers, suggesting both protective and deleterious roles depending on the context. In UVM, higher expression of *SNCA, PINK1,* and *ATP13A2* was associated with improved OS, indicating that these genes might contribute to tumor suppression or adaptive cellular responses that limit tumor aggressiveness.

In contrast, the dual nature of these genes is evident in SKCM, where *SNCA* and *ATP13A2* demonstrated adverse survival correlations, suggesting tumor-promoting roles. These findings emphasize the context-dependent functions of PD-related genes, which may shift between protective and oncogenic depending on the tumor microenvironment, genetic alterations, and immune interactions.

In LGG, the prognostic significance of *PINK1, LRRK2,* and *PRKN/PARK2* was particularly striking, especially in the *TP53* WT subgroup. These findings suggest that the impact of PD-related genes on patient outcomes may rely on functional *TP53* signaling, consistent with their known involvement in DNA damage responses, cell cycle regulation, and apoptotic pathways. Interestingly, the absence of significant associations in *TP53*-mutant contexts further underscores the dependence of these genes on intact p53 functionality, highlighting a potential vulnerability in *TP53*-WT tumors that could be therapeutically exploited.

### TP53-Dependent Mechanisms

The interplay between PD-related gene expression and *TP53* status is evident in several cancers. For instance, *LRRK2*, already mentioned above, and *ATP13A2* demonstrated strong survival associations in *TP53*-WT cancers such as SKCM and LGG but lost significance in *TP53*-mutant contexts. This suggests that these genes may act as co-regulators of p53-mediated pathways, including genomic stability, apoptosis, and stress responses.

The contrasting survival trends for *SNCA* in *TP53*-WT and *TP53*-mutant SKCM further illustrate the complexity of these interactions. In *TP53*-WT tumors, *SNCA* may cooperate with p53 to mitigate tumor progression, while in *TP53*-mutant contexts, it could drive alternative pathways that promote tumor aggressiveness. This duality highlights the need for further mechanistic studies to elucidate the molecular interactions between PD-related genes and p53 signaling.

Interestingly, p53 levels and activity are significantly elevated in the CNS of various cellular and animal models of PD (32), as well as in post-mortem analyses of PD patients’ brains (33). This observation highlights a potential link between cancer and neurodegeneration. All this may suggest that p53 could act as a molecular hub connecting the opposing cellular fates observed in cancer and neurodegeneration, warranting further investigation into its context-dependent roles and potential as a therapeutic target in both conditions (34).

Moreover, this connection may extend beyond PD. In Alzheimer’s disease (AD), the tau protein, encoded by *MAPT*, appears to have a bidirectional relationship with p53, reinforcing the presence of a complex regulatory network that influences both cancer and neurodegeneration (35–36). In general, *TP53* mutations may promote tumor progression by inhibiting apoptosis, whereas, in neurodegeneration, excessive p53 activity may drive neuronal loss.

### Common mechanisms among neurodegenerative processes and cancer

Neurodegenerative processes and cancer have traditionally been viewed as diametrically opposing pathological conditions (37): neurodegeneration is predominantly characterized by the progressive loss of neuronal cells, while cancer is marked by uncontrolled cellular proliferation. This dichotomy suggests fundamentally different underlying mechanisms: neurodegeneration involves processes such as cell death, whereas cancer is driven by cell growth and evasion of programmed cell death (38). However, accumulating evidence reveals that these seemingly distinct conditions share common molecular mechanisms. Both cancer and neurodegeneration involve dysregulation of oxidative stress, inflammation, and genomic instability, the latter being of particular interest (39–40).

Our pathway analysis indicates correlations between the levels of PD-related genes (e.g., *SNCA, LRRK2*, and *PINK1*) and pathways regulating DNA damage response and cell cycle progression, including DNA repair, the G2M checkpoint, DNA replication, and mitotic spindle dynamics. Notably, in brain and skin tumors, these genes negatively correlate with these pathways. In PD, genomic instability and DNA lesions have been identified as potential risk factors for the disease (41). This shared feature suggests that aging-related genomic instability may serve as a genetic modifier predisposing individuals to both cancer and neurodegeneration. Similar findings are relevant to other neurodegenerative disorders, such as AD (42) and Huntington’s disease (43).

Moreover, in diseases like PD and AD, neuronal cells—typically maintained in the G0 phase—can aberrantly reenter the cell cycle, ultimately undergoing mitotic catastrophe (44 and 45). Although the precise mechanisms remain unclear, our pathway analysis highlights correlations with several cell cycle-regulating pathways, underscoring the importance of exploring these shared processes further.

## Conclusions

Overall, our analysis underscores the complex and context-dependent roles of PD-related genes in cancer biology, revealing significant associations with survival, tumor progression, and the interplay with *TP53* status. Importantly, this work is one of the first attempts to bridge the gap between the seemingly opposite fields of neurodegeneration and oncogenesis, shedding light on shared molecular mechanisms, including genomic instability and cell cycle deregulation. The correlation between PD-related genes and pathways regulating DNA repair and mitotic processes suggests that aging-related genomic instability may predispose individuals to both neurodegenerative disorders and cancer. Furthermore, the aberrant cell cycle reentry seen in neurodegenerative diseases, such as PD and AD, mirrors processes observed in certain tumors, reinforcing the notion of a mechanistic continuum between these conditions.

Future research should focus on mechanistic studies to unravel the intricate interactions between these pathways, with the goal of developing treatments that address the overlapping biology of these two conditions.

## Acknowledgment

LC gratefully acknowledges support from Vita-Salute San Raffaele University.

## Authors’ Roles

SF and FM conducted the analysis. LC and MC supervised the project. LC conceived the original idea and wrote the manuscript. All authors reviewed and approved the final manuscript.

## Financial Disclosures of all authors (for the preceding 12 months)

There are no financial conflicts of interest to disclose.

## Data Availability Statement

The data and codes that support the findings of this study are available from the corresponding author upon reasonable request.

